# A glimpse into the DNA virome of the unique “living fossil” *Welwitschia mirabilis*

**DOI:** 10.1101/2022.02.23.481439

**Authors:** Humberto Debat, Nicolás Bejerman

## Abstract

Here, we report the identification and characterization of four novel DNA viruses associated with *Welwitschia mirabilis*. Complete circular virus-like sequences with affinity to *Caulimoviridae* members and geminiviruses were detected and characterized from *Welwitschia mirabilis* genomic data. The two newly *Caulimoviridae*-like viruses have been tentatively named as Welwitschia mirabilis virus 1 and 2 (WMV1-WMV2); whereas the two identified geminiviruses were named as Welwitschia mirabilis associated geminivirus A and B (WMaGVA-WMaGVB). Phylogenetic analysis suggests that WMV1-2 belong to a proposed genus of *Caulimoviridae*-infecting gymnosperms. WMaGVA-B are phylogenetically related with both mastreviruses and capulaviruses and likely represent a distinct evolutionary lineage within geminiviruses.

---

*Welwitschia mirabilis* is a distinctive long-living dwarf tree with only two leaves growing throughout the entire plant life [1]. This plant is a monotypic gymnosperm species that is the lone member of the family *Welwitshchiaceae*, which is endemic to the Namib Desert in both northern Namibia and southern Angola. Welwitschia is usually referred as a “living fossil” because is one of the longest-lived plants on Earth, living up to 3,000 years old [2] and there is a fossil record of this species of 112 My, phylogenetically isolated within the order Gnetales [3]. This exotic and enigmatic plant has been thoroughly studied with a focus in its biological, physiological and ecological peculiarities. Nevertheless, regarding plant health, as of today there has been only one record of a virus linked to this ancient plant: the potyvirus catharanthus mosaic virus (CatMV) which was detected in a Welwitschia plant grown ex situ in a domestic garden in western Australia [4].

A vast number of novel viruses, many of them not inducing any apparent symptoms, have been identified using metagenomic approaches, which revealed our constrained knowledge about the richness and diversity of the plant virosphere [5-7]. In the context of an advancement in the implementation of high-throughput sequencing technologies as a steady tool in virus discovery, it has been stated that those newly identified viruses that are known only from metagenomic data can, should, and have been incorporated into the official classification scheme of the International Committee on Taxonomy of Viruses (ICTV) [8, 9]. In the recent years, the analysis of public data constitutes an emerging source of novel *bona fide* plant viruses, which allows the reliable identification of new viruses in hosts with no previous record of virus infections [10, 11]. In general, plant viruses reported to date based in metagenomic data have been detected analyzing publically available plant transcriptome datasets, resulting mostly in the detection of RNA viruses which could have been inadvertently co-isolated with host RNAs during library preparation [10, 12-15]. WGS DNA datasets have received less attention in virus discovery studies, perhaps given its higher complexity in terms of database size and technical difficulties during processing. However, given their widespread transcription or RNA intermediates it is also possible to detect signatures of DNA viruses from RNAseq datasets, in some cases allowing its characterization and the assembly of complete virus genomes [6, 16-18].

The members belonging to the family *Caulimoviridae* have a double stranded DNA (dsDNA) circular genome with a size ranging between 7.1 kb to 9.8 kb including one to eight open reading frames (ORFs) [19] and are the only plant-infecting viruses known to replicate by reverse transcription [20]. The *Geminiviridae* family is the most numerous family of plant-infecting viruses and its members have genomes comprising one or two single-stranded, circular DNAs (ssDNA) [21].

Here, we analyzed transcriptomic and WGS DNA datasets corresponding to *Welwitschia mirabilis*, publicly available at the NCBI SRA database, which resulted in the identification and assembly of the complete genomic sequences of four novel DNA viruses, the first associated with this “fossil” plant. Two of them are related to *Caulimoviridae* members and were tentatively named as Welwitschia mirabilis virus 1 (WMV1), Welwitschia mirabilis virus 2 (WMV2); whereas the other two novel viruses are related with the geminiviruses and were named as Welwitschia mirabilis associated geminivirus A (WMaGVA) and Welwitschia mirabilis associated geminivirus B (WMaGVB).

The raw data analyzed in this study corresponds to 30 publicly available *Welwitschia mirabilis* RNAseq and WGS DNA NGS libraries (Supplementary Table 1) corresponding to a variety of samples from wild (Namib desert) and ex situ greenhouse plants (China and USA), of different developmental stages (young and old leaves), tissues (meristematic and peripheral meristematic), organs (cones, roots, leaves), from male and female individuals. The RNAseq raw reads from each library were pre-processed by trimming and filtering with the Trimmomatic tool as implemented in http://www.usadellab.org/cms/?page=trimmomatic, the resulting reads were assembled *de novo* with Trinity v2.6.6 release with standard parameters. The WGS DNA dataset was assembled using Unicycler (Version 0.4.8) with standard parameters as implemented in the Galaxy platform https://usegalaxy.org/. *De novo* assembled sequences were subjected to bulk local BLASTX searches (E-value < 1e-5) against a refseq virus protein database available at ftp://ftp.ncbi.nlm.nih.gov/refseq/release/viral/viral.1.protein.faa.gz. The resulting hits were clustered using the CD-HIT tool available at http://weizhong-lab.ucsd.edu/cdhit-web-server/cgi-bin/index.cgi with a sequence identity cut-off of 0.95. The resulting contigs were explored by hand. Our attention was caught on that our best (but substantially low identity) hits corresponded to DNA viruses, more specifically two ca. 2.9 kb sequences from the SRA SRX9189731 corresponding to DNA-Seq of Welwitschia young leaves from a green house in China [22] obtained best significant hits (E-value = 1e-45 and 5e-48, identity = 53.54% and 40.22%) with the replication associated protein of the geminiviruses East African cassava mosaic virus and French bean severe leaf curl virus, respectively. Whereas two ca. 7.6 kb sequences from the SRA SRX4959752 corresponding to RNA-Seq of Welwitschia mature leaves from China [23] obtained best significant hits (E-value = 6e-125 and 5e-107, identity = 29.19% and 36.48%) with the polyprotein encoded by the unclassified *Caulimoviridae-*like virus Pinus nigra virus 1. The tentative virus contigs were extended and curated by iterative mapping of the corresponding libraries raw reads, using a method described by [24]. Essentially, this strategy employs BLAST/nhmmer to extract a subset of reads related to the query contig, use the retrieved reads to extend it and then repeat the process iteratively using as query the extended sequence. Importantly, the sequences extended and polished presented overlapping regions and were subsequently reassembled using Geneious v8.1.9 (Biomatters Ltd.) alignment tool (cost matrix: >93% similarity 5.0/-9.026) into four circular single 2,913 nt (WMaGVA), 2,912 nt (WMaGVB), 7,872 nt (WMV1) and 7,648 nt long (WMV2) sequences. The circular nature of the virus genomes were evidenced by identifying overlapping reads simultaneously covering a continuum of virus reads supporting circularity at all genome positions. Bowtie2 http://bowtie-bio.sourceforge.net/bowtie2/index.shtml was then employed for mean coverage estimation and reads per million (RPMs) calculations. These resulting DNA virus sequences were supported by a total of 2,886 (WMV1), 13,119 (WMV2), 585 (WMaGVA), and 2,241 reads (WMaGVB) (mean coverage = 56.7X, 250.1X, 50.8X and 191.5X, respectively Supplementary Table 1). To further advance in the characterization of these sequences virus ORFs were predicted with ORFfinder (https://www.ncbi.nlm.nih.gov/orffinder/), domains presence and architecture of translated gene products was determined by InterPro (https://www.ebi.ac.uk/interpro/search/sequence-search) and the NCBI Conserved domain database v3.19 (https://www.ncbi.nlm.nih.gov/Structure/cdd/wrpsb.cgi). Further, HHPred and HHBlits as implemented in https://toolkit.tuebingen.mpg.de/#/tools/ were used to complement annotation of divergent predicted proteins by hidden Markov models. In order to infer phylogenetic relationships, the aa sequences of the WMV1, WMV2, WMaGVA and WMaGVB selected proteins (RT-RNase H domain for caulimovirids and CP and Rep for the geminiviruses), as well as the WMaGVA and WMaGVB complete genomic sequences, were aligned with the corresponding sequences of caulimovirids and geminiviruses described in Supplementary Table 2, using MAFTT 7 https://mafft.cbrc.jp/alignment/software/ (BLOSUM62 scoring matrix) using as best-fit algorithm G-INS-i (RT, CP and Rep) and E-INS-i (complete genomic sequences). The aligned sequences were subsequently used as input for FastTree 2.1.9 (http://www.microbesonline.org/fasttree/) ML phylogenetic trees computing local support values with the Shimodaira-Hasegawa test (SH) and 1,000 tree resamples or MegaX [25] where 1000 bootstrap replicates were used to construct the ML trees.

The assembled sequences corresponding to the complete genome sequence of WMV1, WMV2, WMaGVA and WMaGVB were deposited in GenBank under accession numbers BK061147, BK061148, BK061149 and BK061150, respectively.

The genome of WMV1 and WMV2 were determined to be 7,872 and 7,648 nt in size, respectively, and they have one single large open reading frame (ORF) coding a 2,264 and 2,111 amino acids (aa) polyprotein, respectively (Fig. 1A). The WMV1 encoded polyproteins have several conserved domains: movement protein (aa 63 to 177 coordinates), reverse transcriptase (aa 1,419 to 1,667), and RT-RNase H (aa 1,689 to 1,810); whereas a movement protein (aa 57-188), Peptidase A3 (aa 932 to 1,080), reverse transcriptase (aa 1,124 to 1,302), and RNase H (aa 1,399 to 1,559) conserved domains were identified in the WMV2 polyprotein. Similar domains were described in the identified *Caulimoviridae* virus and best hit Pinus Nigra virus 1 (PNV1) [26] and other caulimovirids [19].

**Fig. 1.**
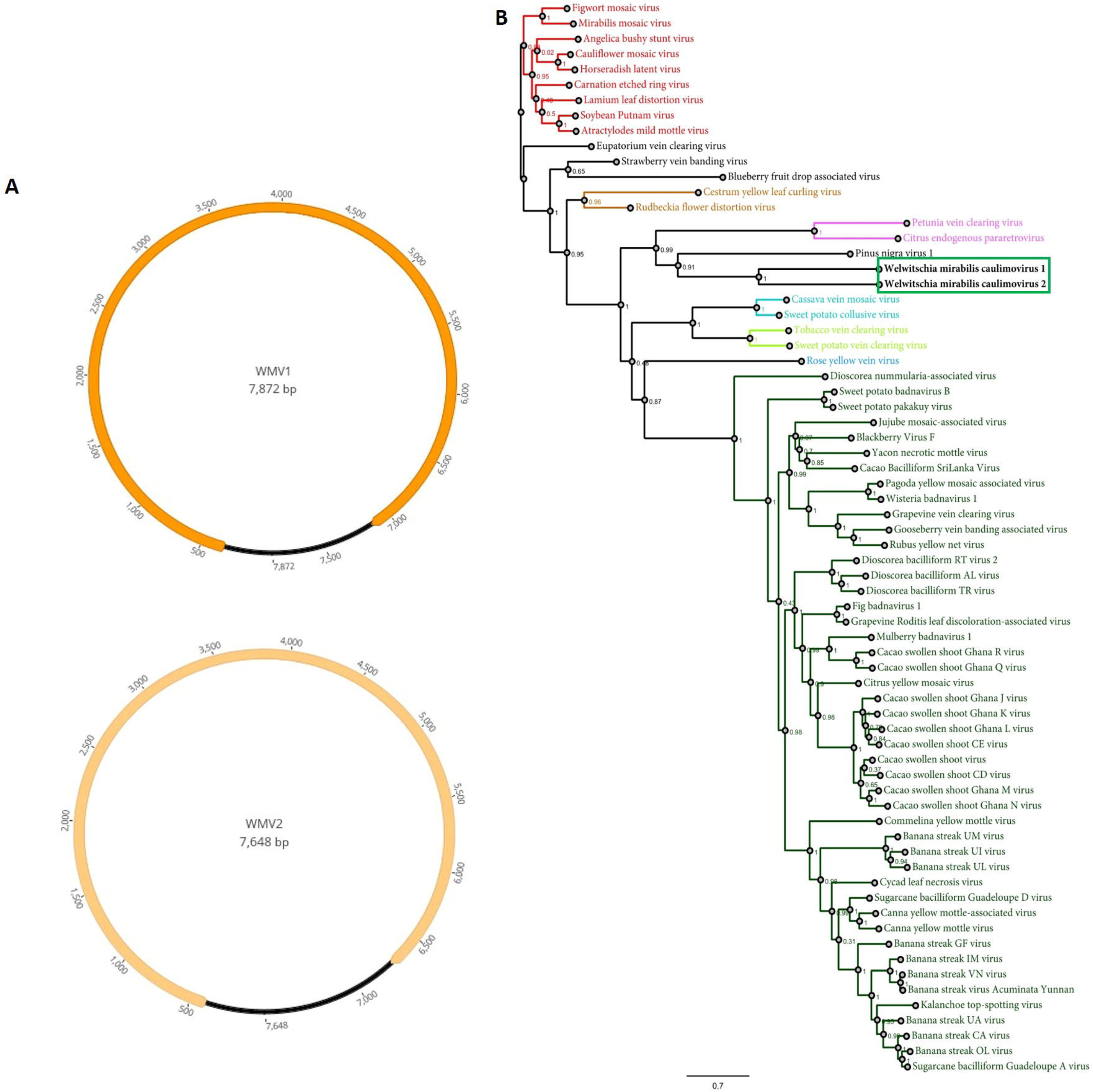
(A) Genomic organization of Welwitschia mirabilis virus 1 (WMV1) and Welwitschia mirabilis virus 2 (WMV2). (B) Maximum likelihood phylogenetic tree analyzing the WMV1 and WMV2 and representative caulimovirids RT-RNase H domain amino acid sequences. The bar below each tree represents substitutions per site. Accession numbers of every virus used to construct phylogenetic trees are listed in Supplementary Table 2.

The WMV1 and WMV2 polyprotein shared just a 28.3% aa identity and their polyproteins shared the highest identity (36% and 29%, respectively) with that one encoded by PNV1; whereas the WMV1 and WMV2 RT-RNase H domain, which is currently used for the demarcation of species in the family *Caulimoviridae* [27] shared a 48.2% aa identity between them, and a highest identity of 54.14% and 52.78%, respectively, with the similar domain of PNVI1. Therefore, the substantially low genetic identity of WMV1 and WMV2 between them and with other *caulimoviridae* members suggests that these two viruses should be classified as members of two new species.

In order to determine the phylogenetic relationships of WMV1 and WMV2 with other caulimovirids to unravel its possible evolutionary history, maximum likelihood (ML) phylogenetic analyses were carried out using aa sequences of the RT-RNase H domain. ML phylogenetic trees were constructed using G-INS-i as the best-fit model. Phylogenetic analysis showed that WMV1 and WMV2 clustered together with PNV1 (Fig. 1B). PNVI has been proposed to represent a new genus of *Caulimoviridae*-infecting gymnosperms [26]. Thus, WMV1 and WMV2 should belong, along with PNV1, to this recently proposed novel genus within the *Caulimoviridae* family, which could be named as *Gymnovirus*, because all these viruses have been identified in gymnosperms and are phylogenetically related, so they likely share a common evolutionary history consistent with infection of gymnosperm hosts, an understudied and neglected clade of spermatophytes in plant virology.

The WMaGVA and WMaGVB genomes are monopartite and were determined to be 2,913 and 2,912 nt in length, respectively. Their circular DNA contains a nonanucleotide sequence TATATTAAC, that indicates the probable origin of virion-strand and DNA replication, slightly different than the nonanucleotide sequence TAATATTAC, which is found in most of the geminiviruses described so far [28]. Five putative proteins were identified in the circular ssDNA genome of WMaGVA and WMaGVB. The inferred genome organization of these viruses has three virion-sense (V1, V2 and V3) and two complementary-sense (C1 and C1:C2) genes with two intergenic regions (LIR and SIR (Fig. 2A). Similar to Curtoviruses [21], the two novel viruses have three ORFs in the virion-sense strand with the same genomic architecture. Whereas the number and genomic location of the ORFs in the complementary-sense of WMaGVA and WMaGVB is similar to that described for Becurtoviruses and Mastreviruses [21]. The genes on sense and complementary strands are separated by two intergenic regions, a relatively short one (SIR) that separates the genes V1 and C1:C2, and a long one (LIR) that separates the genes C1 and V3 (Fig. 2A). A LIR and a SIR is present in several genera belonging to the *Geminiviridae* family [21, 28].

**Fig. 2.**
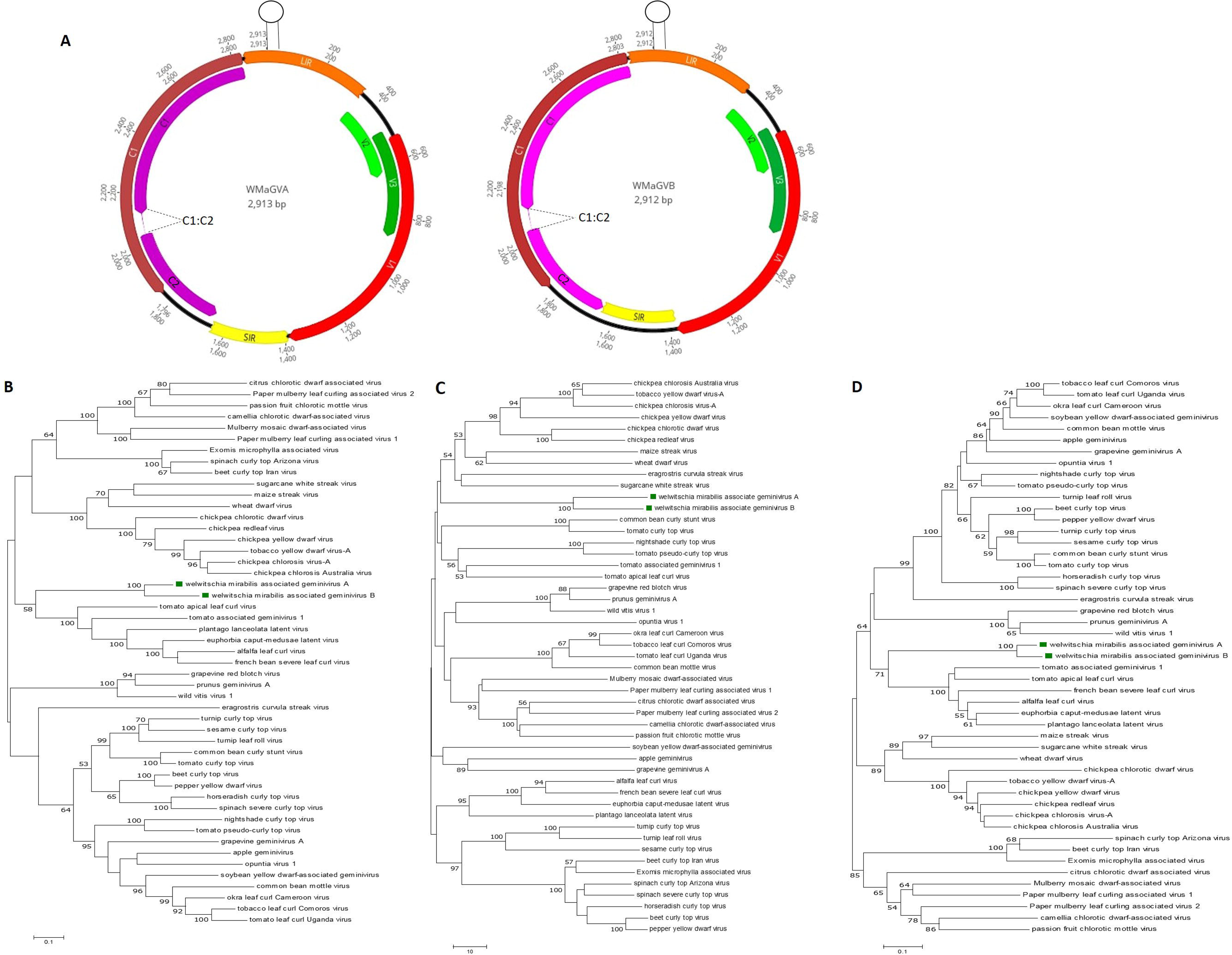
(A) Genomic organization of Welwitschia mirabilis associated geminivirus A (WMaGVA) and Welwitschia mirabilis associated geminivirus B (WMaGVB). Maximum likelihood phylogenetic tree analyzing the WMaGVA and WMaGVB and representative geminiviruses complete genome nucleotide sequences (B), as well as CP (C) and Rep (D) amino acid sequences. The bar below each tree represents substitutions per site. Accession numbers of every virus used to construct phylogenetic trees are listed in Supplementary Table 2.

A BLASTp search analysis of the predicted proteins of WMaGVA and WMaGVB showed that two of their five encoded proteins, encoded by V2 and V3 genes, had no significant sequence similarity to any others sequences in the database (Table 1). However, one transmembrane domain was identified in the WMaGVA and WMaGVB V2 ORF encoded protein; transmembrane domains were also predicted in the putative movement protein (MP) of several geminiviruses [29]. Furthermore, the ORF V2 is immediately upstream of the capsid protein (CP) (Fig. 2A), where the putative movement proteins are usually located in monopartite members of *Geminiviridae* family [28]; thus, V2 gene likely encodes a MP. No conserved motif or domains were identified in the product encoded by the V3 gene. The WMaGVA and WMaGVB ORF V1 (Fig.1) is similar to the CP encoded by mastreviruses. The replication-associated protein A (RepA) is encoded by the C1 gene; while the replicase protein (Rep) is expressed from C1 and C2 genes potentially by transcript splicing and contains the conserved motifs described for this protein in other geminiviruses [30]. A similar splicing of C1:C2 was also described for becurtovirues, capulaviruses, grabloviruses and mastreviruses [21]. Unlike the V1 ORF encoded protein, the translation product of ORF C1:C2 is similar to the Rep encoded by capulaviruses. In summary, the V1 gene encoded protein was identified as the putative CP, while the C1:C2 encoded protein was identified as the putative Rep protein, while V1 likely encodes the putative movement protein and C1 encodes the RepA protein.

**Table 1.**
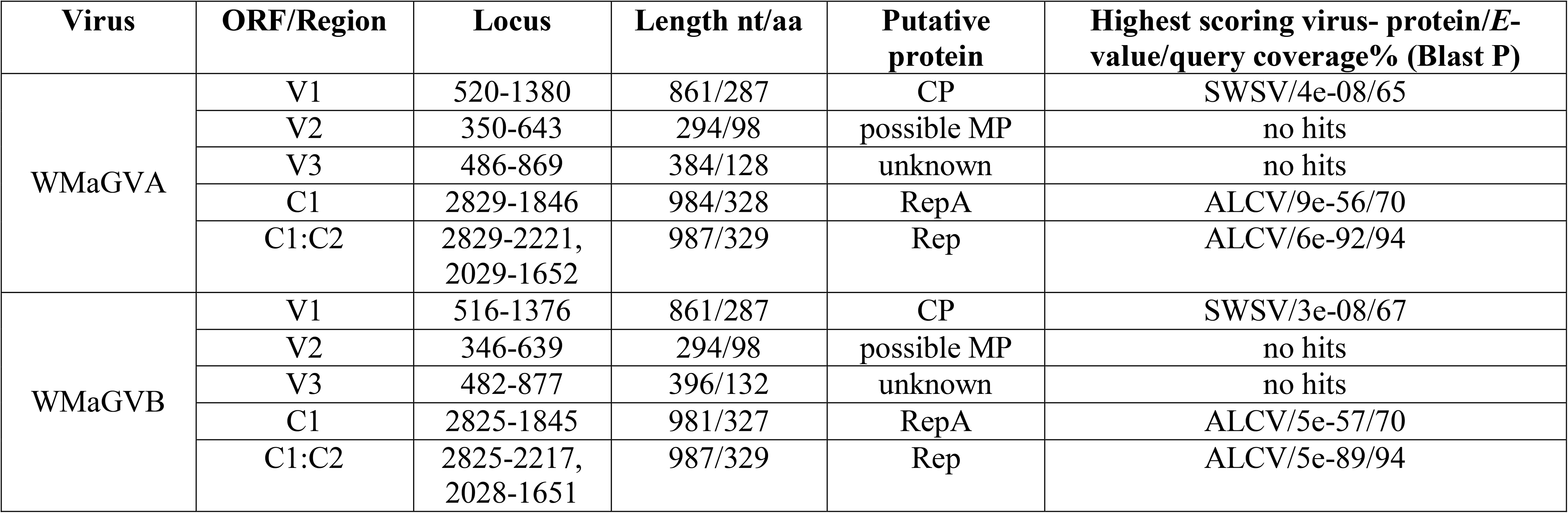
Genomic organization of Welwitschia mirabilis associated geminivirus A (WMaGVA) and Welwitschia mirabilis associated geminivirus B (WMaGVB)

Genome-wide pairwise comparisons using the Sequence Demarcation Tool (SDT v1.2) [31] indicated that WMaGVA and WMaGVB share 77.2% identity between each other, and these viruses shared a maximum of 60.5% identity with all of other known geminiviruses (Supplementary Fig. 1A). In pairwise comparisons, the WMaGVA and WMaGVB CP aa sequences shared 84.32% identity between each other, and a highest identity (30.71%) with the mastrevirus sugarcane white streak virus (SWSV) (Supplementary Fig. 1B); whereas WMaGVA and WMaGVB Rep protein sequences shared 87.16% identity between each other, and a highest identity (47.3%) with the capulavirus alfalfa leaf curl virus (Supplementary Fig. 1C). These results showed that these viruses are distantly related to other geminiviruses. Therefore, the deeply low genetic identity of WMaGVA and WMaGVB with other geminiviruses suggests that these two viruses should be classified as members of a new species that could belong to a novel genus within family *Geminiviridae*, which could be named as *Welwitschiavirus*.

Phylogenetic analysis of the WMaGVA and WMaGVB sequences together with the full-genome (or DNA-A sequence) of other known geminiviruses, using E-INS-i as the best-fit model, indicated that these two viruses formed a distinct clade that clustered with a group formed by the capulaviruses members as well the unassigned geminiviruses tomato apical leaf curl virus and tomato associated geminivirus 1 (Fig.2B). Phylogenetic analyses carried out using the aa sequences of the CP and Rep genes with G-INS-i as the best-fit model showed that WMaGVA and WMaGVB CP formed a distinct clade that clustered with the a group formed by mastreviruses (Fig. 2C); whereas the Rep protein based tree showed that WMaGVA and WMaGVB Rep protein formed a distinct clade that clustered with a group formed by capulaviruses members as well the unassigned geminiviruses tomato apical leaf curl virus and tomato associated geminivirus 1 (Fig. 2D), which further supports the results obtained when both the CP and Rep amino acid sequences were analyzed using SDT. Phylogenetic analysis placed WMaGVA and, WMaGVB in a unique taxon within geminiviruses; therefore it could be concluded that this virus represents and evolutionary distinct geminivirus member. Furthermore, the different evolutionary relationships of the Rep and CP proteins likely support a modular organization and a different evolutionary history of the virion-sense and complementary-sense frame of the WMaGVA and WMaGVB genomes. Several geminiviruses with chimeric genomes architecture and phylogeny, such as WMaGVA and WMaGVB were identified [32-36]. Moreover, recently, it was shown that the CP sequences of geminiviruses are likely co-diverging with their insect vector [37]. The distinct phylogenetic cluster formed by WMaGVA and WMaGVB indicates that these viruses are likely transmitted by a vector that has not yet been described for geminiviruses, which in turn should involve an insect adapted to the peculiar Namib coastal desert, one of the driest and oldest desert regions of the world.

It is apparent that the WMaGVA and WMaGVB are chimeric virus encoding Rep and CP with differential evolutionary histories because their CP and Rep phylogenies are not congruent; thus, it is likely that these viruses are a recombinant of mastre-like and capula-like ancestral viruses. However, when recombination analysis of full-length genomes were performed using RDP, GENECONV, MaxChi, BootScan, 3Seq, Chimera and SiScan statistics methods implemented in the RDP5 software [38], no recombination events were detected in the WMaGVA and WMaGVB genome sequences. This result could be explained by the fact that the non-recombinant descendants of WMaGVA and WMaGVB parents have not been described yet or no longer exist.

WMaGVA and WMaGVB display a geminivirus-like genomic size and organization, and encodes a CP that is most closely related to that of mastreviruses, and a Rep protein that is most closely related to that of capulaviruses. Thus, these viruses represent an evolutionary distinct lineage within the geminiviruses and therefore they are unique novel members that may represent a new genus within the *Geminiviridae* family.

Given the difficulties associated to identifying a proper host based only on metagenomics analyses of NGS data, we extended our search using as query the genomic sequences of the characterized viruses from the initial two libraries to the assemblies generated for the additional 28 experiments. Interestingly we found blastn hits ranging from 96-100% identity in 20% of the assessed libraries for WMV1 and WMV2, and blastn hits ranging from 93-100% identity in 90% of the libraries for WMaGVA and WMaGVB. We confirmed theses detections by assessing the raw data by read mapping using Bowtie2 with standard parameters against the consensus genomes (Supplementary Table 1) which was consistent with the detection of these viruses in multiple independent samples from independent studies obtained from wild and ex situ plants from the Namib Desert, China and USA, from diverse tissues, organs and developmental stages. These results provide further robust evidence on the association of these viruses with Welwitschia and suggest that the caulimo and geminiviruses reported here appear to be prevalent in this host.

In summary, the analysis of public SRA data is a valuable source to detect potential novel DNA plant viruses. Using this approach, we explored a glimpse into the DNA virome of the fossil plant *Welwitschia mirabilis* which resulted in the identification and reconstruction of the complete genome of two novel geminiviruses and two novel caulimovirids associated with *Welwitschia mirabilis*. This is the first report of caulimo and geminiviruses infecting this ancient gymnosperm plant worldwide. Future studies should assess many intriguing aspects on the biology and ecology of these viruses linked with such a peculiar host in a hostile and restricted habitat, their prevalence, their vectors, and some clues about their potential impact on their host.

## Acknowledgments

We would like to express a sincere gratitude to the generators of the underlying data used for this work. By following open access practices and supporting accessible raw sequence data in public repositories available to the research community, they have promoted the generation of new knowledge and ideas.

## Statements and Declarations

### Author Contributions

Humberto Debat and Nicolas Bejerman contributed to the study conception and design, data analysis. The manuscript was written by both authors, who commented and reviewed it. All authors read and approved the final manuscript.

### Data Availability

The virus genome sequences are deposited in database Genbank under accession numbers BK061147, BK061148, BK061149 and BK061150.

### Ethical Approval

This article does not contain any studies with human participants or animals performed by any of the authors.

