## Supplementary material for "A glimpse into the DNA virome of the unique “living fossil” *Welwitschia mirabilis*": Supp Fig. 1

**Supplementary Figure 1**. Pairwise identity matrix of Welwitschia mirabilis associated geminivirus A (WMaGVA) and Welwitschia mirabilis associated geminivirus B (WMaGVB), and representative geminiviruses complete genomes (A) nucleotide sequences, as well as CP (B) and Rep (C) amino acid sequences generated using the SDT v1.2 software (Muhire et al., 2014). Accession numbers of every virus are listed in Supplementary Table 2.

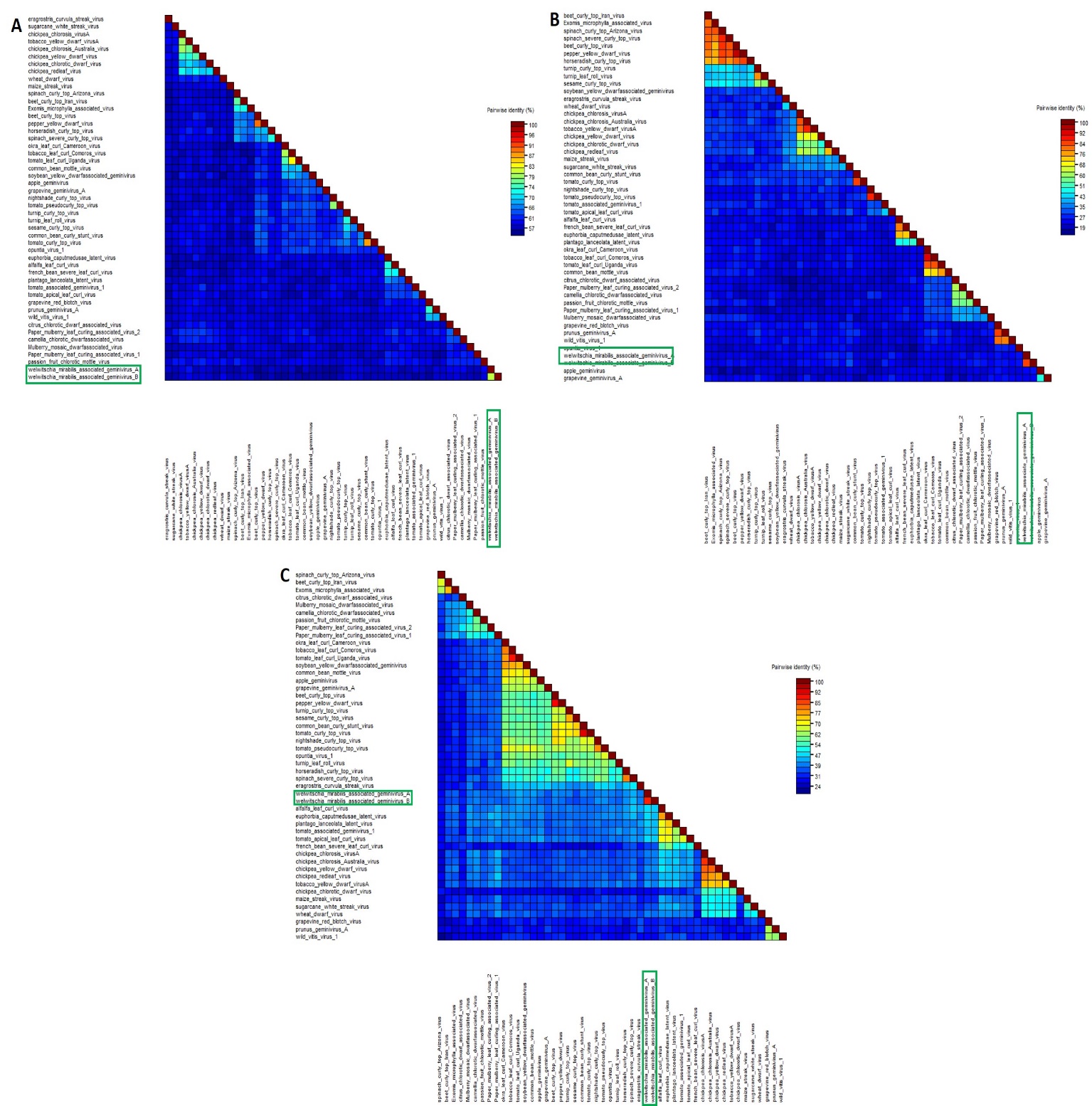
