## Supplementary Table 1 for "A glimpse into the DNA virome of the unique “living fossil” *Welwitschia mirabilis*"

**Supplementary Table 2.** Accession numbers of every virus isolate used to construct the phylogenetic trees.

| **Virus name** | **Accession number** |
| --- | --- |
| Okra leaf curl Cameroon virus | HE793426 |
| Tobacco leaf curl Comoros virus | AM701762 |
| Tomato leaf curl Uganda virus | MN38114 |
| Chickpea chlorosis Australia virus | JN989418 |
| Chickpea chlorosis virus-A | JN989414 |
| Chickpea chlorotic dwarf virus | KF176553 |
| Tobacco yellow dwarf virus-A | KC172702 |
| Alfalfa leaf curl virus | KX574859 |
| Horseradish curly top virus | U49907 |
| Beet curly top Iran virus | EU273818 |
| Exomis microphylla associated virus | MG001960 |
| common bean mottle virus | KX011473 |
| French bean severe leaf curl virus | JX094280 |
| Plantago lanceolata latent virus | KT214389 |
| Spinach curly top Arizona virus | HQ443515 |
| Euphorbia caput-medusae latent virus | KT214384 |
| Beet curly top virus | M24597 |
| Spinach severe curly top virus | GU734126 |
| Eragrostris curvula streak virus | FJ665631 |
| Grapevine red blotch virus | KC427996 |
| Citrus chlorotic dwarf associated virus | KF561253 |
| Prunus geminivirus A | MF504034 |
| Wild vitis virus 1 | MG976084 |
| Tomato associated geminivirus 1 | MF072688 |
| Maize streak virus | AF007881 |
| Sugarcane white streak virus | KJ187749 |
| Wheat dwarf virus | DQ868525 |
| Chickpea redleaf virus | GU256532 |
| Chickpea yellow dwarf virus | KM377674 |
| Mulberry mosaic dwarf-associated virus | MN240483 |
| Camellia chlorotic dwarf-associated virus | MK613869 |
| Common bean curly stunt virus | MK673513 |
| Grapevine geminivirus A | KX570609 |
| Apple geminivirus | MT107729 |
| turnip curly top virus-A | KX533468 |
| turnip curly top virus-B | GU456685 |
| sesame curly top virus | MH595454 |
| turnip leaf roll virus | KT388078 |
| tomato pseudo-curly top virus | X84735 |
| nightshade curly top virus | MT521865 |
| Passion fruit chlorotic mottle virus | MG696802 |
| Opuntia virus 1-A | MN100038 |
| Opuntia virus 1-B | MN100009 |
| Opuntia virus 1-C | MN100035 |
| Opuntia virus 1-D | MN099983 |
| Tomato apical leaf curl virus | MH491195 |
| Paper mulberry leaf curling associated virus 1 | MN595124 |
| Paper mulberry leaf curling associated virus 2 | MN595127 |
| pepper yellow dwarf virus | EU921828 |
| soybean yellow dwarf-associated geminivirus | MT221447 |
| tomato curly top virus | AB935398 |
| Temperate fruit decay-associated virus-MFB13 | KJ955448 |
| Temperate fruit decay-associated virus-MFB4 | KR134342 |
| Temperate fruit decay-associated virus-MFBpe1 | KJ955450 |
| Temperate fruit decay-associated virus-MFBpe23 | KJ955447 |
