## Supplementary Table 2 for "A glimpse into the DNA virome of the unique “living fossil” *Welwitschia mirabilis*"

**Supplementary Table 1.** NCBI publicly available datasets used for virus discovery. Total virus reads and per million are based on mapping results using Bowtie.2 with standard parameters against the genome consensus sequences of Welwitschia mirabilis virus 1 and 2 (WMV1, WMV2) and Welwitschia mirabilis associated geminivirus A and B (WMaGVA, WMaGVB).

| **Run** | **BioProject** | **BioSample** | **Experiment** | **Total library reads** | **WMaGVA reads** | **WMaGVA RPM** | **WMaGVB reads** | **WMaGVB RPM** | **WMV1 reads** | **WMV1 RPM** | **WMV2 reads** | **WMV2 RPM** |
| --- | --- | --- | --- | --- | --- | --- | --- | --- | --- | --- | --- | --- |
| SRR8138861 | PRJNA503117 | SAMN10352752 | SRX4959752 | 55427070 | 1,354 | **24,4** | 1,707 | **30,8** | 2,886 | **52,1** | 13,119 | **236,7** |
| SRR5894351 | PRJNA396655 | SAMN07430725 | SRX3059925 | 53682066 | 654 | **12,2** | 849 | **15,8** | 28 | **0,5** | 49 | **0,9** |
| ERR845262 | PRJEB9038 | SAMEA3333553 | ERX926173 | 3748384 | 27 | **7,2** | 25 | **6,7** | 9 | **2,4** | 13 | **3,5** |
| ERR364404 | PRJEB4921 | SAMEA2241918 | ERX337173 | 22416158 | 913 | **40,7** | 530 | **23,6** | 1 | **0,0** | 42 | **1,9** |
| ERR3588923 | PRJEB18807 | SAMEA44449918 | ERX3584808 | 3748384 | 22 | **5,9** | 28 | **7,5** | 6 | **1,6** | 14 | **3,7** |
| SRR12710831 | PRJNA665158 | SAMN16244547 | SRX9189731 | 74248360 | 585 | **7,9** | 2,241 | **30,2** | 331 | **4,5** | 137 | **1,8** |
| SRR13249257 | PRJNA680422 | SAMN16953877 | SRX9680955 | 46777930 | 443 | **9,5** | 332 | **7,1** | 6 | **0,1** | 3 | **0,1** |
| SRR13249258 | PRJNA680422 | SAMN16953877 | SRX9680954 | 40746078 | 319 | **7,8** | 246 | **6,0** | 9 | **0,2** | 9 | **0,2** |
| SRR13249264 | PRJNA680422 | SAMN16953877 | SRX9680948 | 41077282 | 293 | **7,1** | 209 | **5,1** | 4 | **0,1** | 4 | **0,1** |
| SRR13249259 | PRJNA680422 | SAMN16953877 | SRX9680953 | 46289762 | 190 | **4,1** | 148 | **3,2** | 0 | **0,0** | 7 | **0,2** |
| SRR13249260 | PRJNA680422 | SAMN16953877 | SRX9680952 | 42813418 | 475 | **11,1** | 383 | **8,9** | 0 | **0,0** | 2 | **0,0** |
| SRR13249272 | PRJNA680422 | SAMN16953877 | SRX9680940 | 42662914 | 260 | **6,1** | 192 | **4,5** | 0 | **0,0** | 0 | **0,0** |
| SRR13249261 | PRJNA680422 | SAMN16953877 | SRX9680951 | 49031728 | 45 | **0,9** | 29 | **0,6** | 0 | **0,0** | 1 | **0,0** |
| SRR13249262 | PRJNA680422 | SAMN16953877 | SRX9680950 | 49614190 | 69 | **1,4** | 84 | **1,7** | 0 | **0,0** | 5 | **0,1** |
| SRR13249263 | PRJNA680422 | SAMN16953877 | SRX9680949 | 53959530 | 147 | **2,7** | 171 | **3,2** | 1 | **0,0** | 3 | **0,1** |
| SRR13249265 | PRJNA680422 | SAMN16953877 | SRX9680947 | 41841924 | 77 | **1,8** | 96 | **2,3** | 0 | **0,0** | 9 | **0,2** |
| SRR13249266 | PRJNA680422 | SAMN16953877 | SRX9680946 | 43181020 | 109 | **2,5** | 108 | **2,5** | 0 | **0,0** | 5 | **0,1** |
| SRR13249267 | PRJNA680422 | SAMN16953877 | SRX9680945 | 39928056 | 143 | **3,6** | 152 | **3,8** | 1 | **0,0** | 4 | **0,1** |
| SRR13249268 | PRJNA680422 | SAMN16953877 | SRX9680944 | 43677554 | 165 | **3,8** | 45 | **1,0** | 3 | **0,1** | 2 | **0,0** |
| SRR13249269 | PRJNA680422 | SAMN16953877 | SRX9680943 | 50379824 | 179 | **3,6** | 39 | **0,8** | 0 | **0,0** | 2 | **0,0** |
| SRR13249270 | PRJNA680422 | SAMN16953877 | SRX9680942 | 46830048 | 290 | **6,2** | 67 | **1,4** | 10 | **0,2** | 8 | **0,2** |
| SRR13249271 | PRJNA680422 | SAMN16953877 | SRX9680941 | 27230766 | 307 | **11,3** | 360 | **13,2** | 10,045 | **368,9** | 2,710 | **99,5** |
| SRR13249273 | PRJNA680422 | SAMN16953877 | SRX9680939 | 38954524 | 783 | **20,1** | 1,774 | **45,5** | 11,683 | **299,9** | 1,925 | **49,4** |
| SRR13249274 | PRJNA680422 | SAMN16953877 | SRX9680938 | 37097070 | 611 | **16,5** | 1,661 | **44,8** | 11,463 | **309,0** | 1,536 | **41,4** |
| SRR13249275 | PRJNA680422 | SAMN16953877 | SRX9680937 | 49676946 | 188 | **3,8** | 188 | **3,8** | 10 | **0,2** | 2 | **0,0** |
| SRR13249276 | PRJNA680422 | SAMN16953877 | SRX9680936 | 42516988 | 151 | **3,6** | 161 | **3,8** | 4 | **0,1** | 17 | **0,4** |
| SRR13249277 | PRJNA680422 | SAMN16953877 | SRX9680935 | 43324542 | 135 | **3,1** | 118 | **2,7** | 12 | **0,3** | 14 | **0,3** |
| SRR13249278 | PRJNA680422 | SAMN16953877 | SRX9680934 | 37392338 | 151 | **4,0** | 147 | **3,9** | 16 | **0,4** | 2 | **0,1** |
| SRR13249279 | PRJNA680422 | SAMN16953877 | SRX9680933 | 39063664 | 177 | **4,5** | 144 | **3,7** | 0 | **0,0** | 1 | **0,0** |
| SRR13249280 | PRJNA680422 | SAMN16953877 | SRX9680932 | 44260368 | 315 | **7,1** | 297 | **6,7** | 6 | **0,1** | 4 | **0,1** |
